# compare_genomes: a comparative genomics workflow to streamline the analysis of evolutionary divergence across genomes

**DOI:** 10.1101/2023.03.16.533049

**Authors:** Jefferson Paril, Tannaz Zare, Alexandre Fournier-Level

## Abstract

**Summary:** The dawn of cost-effective genome assembly is enabling deep comparative genomics to address fundamental evolutionary questions by comparing the genomes of multiple species. However, comparative genomics analyses often deploy multiple, often purpose-built frameworks, limiting their transferability and replicability. Here, we developed compare_genomes, a transferable and extensible comparative genomics workflow package which streamlines the identification of orthologous families within and across genomes and tests for the presence of several mechanisms of evolution (gene family expansion or contraction and substitution rates within protein-coding sequences).

**Availability and Implementation:** The workflow is available for Linux, written as a Nextflow workflow which calls established genomics and phylogenetics tools to streamline the analysis and visualisation of genome divergence. This workflow is freely available at https://github.com/jeffersonfparil/compare_genomes, distributed under the GNU General Public License version 3 (GPLv3).

**Contact:** Corresponding author: Jeff Paril. Queries and issues regarding the implementation can be submitted on the issue page of the github repository: https://github.com/jeffersonfparil/compare_genomes/issues.

**Supplementary information:** [Synonymous to non-synonymous (Ka/Ks) nucleotide substitution ratio plots for the example data set are found in the github repository](https://github.com/jeffersonfparil/compare_genomes/tree/main/misc/KaKs_ratio_plots).

## Introduction

The genomes of related organisms represent records of evolutionary histories along the tree of life. We can infer the drivers of their evolution by analysing the signatures of genome evolution during polyploidisation events, gene duplication or loss, or selection of adaptive mutations between species. This comparative genomics framework scrutinises alternative evolutionary histories across different species within and between clades which will offer insights into how the spatio-temporal dynamics and interactions between the biosphere and the environment prefer one group of biological solutions to fitness over others.

The increasing availability of affordable high-throughput DNA sequencing has allowed the assembly of the genomes of multiple species beyond the early set of model species towards species of specific biological relevance. This enabled the use of deep comparative genomics to answer fundamental evolutionary questions using multiple species within and across clades. However, the pipelines for these analyses are often study-specific and rarely described with enough details to be fully reproducible. This is a major impediment to the generalisation or meta-analysis of the results and hinders the transfer of these workflows. Hence, the bioinformatics community would benefit from a unified but open-source and portable comparative genomics workflow.

Web-based comparative genomics workflows exist (eg. PLAZA; Van Bel et al, 2022). However, the centralised design limits potential extension and usage is inherently limited to the computational resources provided, making it unsuitable to high throughput or high bandwidth usage. Various portable frameworks have been developed to run comparative genomics pipelines in a reproducible way (e.g. Snakemake by Mölder et al, 2021 and Nextflow by Di Tommaso et al, 2017). These tools work synergistically with package and environment management systems such as Docker (https://www.docker.com/) or Conda (https://docs.conda.io/en/latest/). These were used to generate genome assemblies and annotations, sequence alignments, variant callings, and transcriptomic data analyses. However, there is a noticeable lack of fully transparent and easily transferable comparative genomics analysis workflows. In this application note, we address this gap with compare_genomes, a comparative genomics workflow built under the Nextflow framework with packages managed by Conda.

### Features and implementation

The compare_genomes workflow consists of nine analysis steps under the default setup (Figure 1: left panel). These steps are:

**Figure 1:**
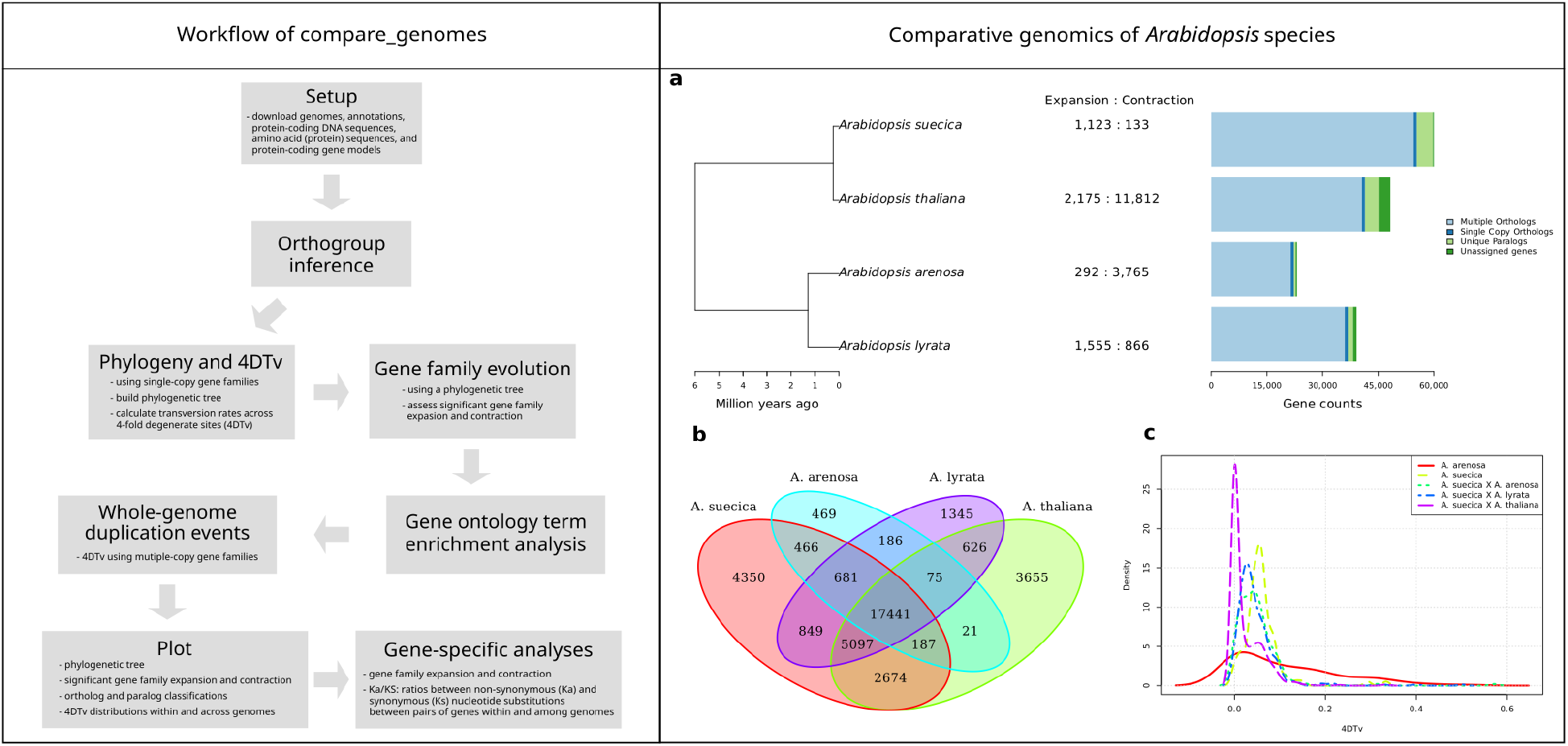
**Left panel:** The steps performed by the compare_genomes comparative genomics workflow. **Right panel:** A sample output plot generated by the compare_genomes workflow using four *Arabidopsis* species.

1. Download of the user-defined genome datasets (i.e. genome sequences, annotations, coding DNA sequences or CDS, protein sequences, protein-coding gene models, corresponding gene ontology terms, and protein sequences of specific genes of interest).
2. Identification of orthogroups using OrthoFinder (Emms and Kelly 2019) and within genome gene family for each orthogroup using HMMER3 (Mistry et al, 2013) and Panther HMMs (protein-coding gene family models; Thomas et al, 2022).
3. Inference of phylogenomic trees for each orthogroup with IQ-TREE 2 (Minh et al, 2020) using CDS alignments generated by MACSE (Ranwez et al, 2011) and the most likely nucleotide substitution model inferred by ModelFinder (Kalyaanamoorthy et al, 2017).
4. Inference of the rate of molecular evolution based on transversion rates among four-fold degenerate sites (4DTv) in single-copy genes between each pair of species using custom Julia scripts.
5. Assessment of whole-genome duplication events using 4DTv computed from multi-copy gene families.
6. Test of significant gene family expansion or contraction across genomes using CAFE (version 5; De Bie et al, 2006).
7. Gene ontology (GO) term enrichment analysis for significantly expanded gene families using the Panther GO API (Mi et al, 2019).
8. Visualisation of a summary of the results (i.e. Figure 1: right panel for a sample output).
9. Analysis of user-defined gene(s) of interest, i.e. gene family expansion/contraction analyses with CAFE, and estimation of non-synonymous to synonymous nucleotide substitution rates (Ka/Ks) using KaKs_Calculator 2.0 (Wang et al, 2010) and custom R script.

Compare_genomes was implemented using Nextflow because of the ease of integrating new and existing Linux-based bioinformatics pipelines. Analysis steps can be easily added or modified, for example adding a GO term enrichment analysis for the significantly contracted families, or substitute MACSE for another multiple sequence alignment tool.

### Usage and example

A detailed user manual describing how to install, set up, and run the workflow is presented in the README page of the compare_genomes project repository: https://github.com/jeffersonfparil/compare_genomes. We included a tutorial and dataset analysing four Pseudomonas spp. species. The compare_genomes workflow was initially conceived to compare the newly released, high-quality reference genome of annual ryegrass (*Lolium rigidum*; an important weed in winter cropping) to related grass species, including the pasture crop perennial ryegrass (*Lolium perenne*). This analysis showed significant expansion of herbicide resistance-related gene families, including detoxification genes in the noxious weed annual ryegrass (Paril et al, 2022).

To illustrate the performance of our pipeline, we compared the genomes of four well characterised *Arabidopsis* species to test if it is able to recapture the evolutionary patterns expected in these species. The summary of the output of this example is shown in Figure 1 right panel. It shows that the phylogenetic relationship between species was accurately recapitulated as expected from the results of Novikova and colleagues in 2016. This summary figure also shows the differences in patterns of gene family contraction and expansion. *Arabidopsis suecica*, an allopolyploid hybrid of A. *thaliana* and A. *arenosa* (Novikova et al, 2017; Burns et al, 2021), experienced gene family expansion. A similar expansion was observed in A. *lyrata*, an outbreeder which diverged from A. *thaliana* around 5 million years ago (Schmickl et al, 2010). Similarities in gene family composition between species are presented as a Venn diagram. Finally, the presence of whole-genome duplication events was assessed using 4DTv density plot. This analysis recaptured the recent polyploidisation event of the A. *arenosa* sub-genome within the A. *suecica* allopolyploid genome detected by Novikova and colleagues in 2017. The Ka/Ks ratio analyses presented in the supplementary information show evidence of selection across several 15-bp windows in the GSTU13 (glutathione transferase - tau class; size of the window can be modified).

## Conclusion

We developed compare_genomes, a transferable and extendible comparative genomics workflow built using the Nextflow framework and Conda package management system. It provides a wieldy pipeline to test for non-random evolutionary patterns which can be mapped to evolutionary processes to help identify the molecular basis of specific features or remarkable biological properties of the species analysed. Additionally, it provides a template which other comparative genomics pipelines can be built or patterned upon for improved reproducibility.

## Acknowledgements

We thank the Department of Agriculture Fisheries and Forestry of Australia together with the Grains Research and Development Corporation (Grant ID: 4-FY9JQPE), Australian Research Data Commons (Grant ID: DP727), Commonwealth Scientific and Industrial Research Organisation for the funding, and the University of Melbourne for hosting this study.

## Supplementary Information

**Supplementary Figure 1:**
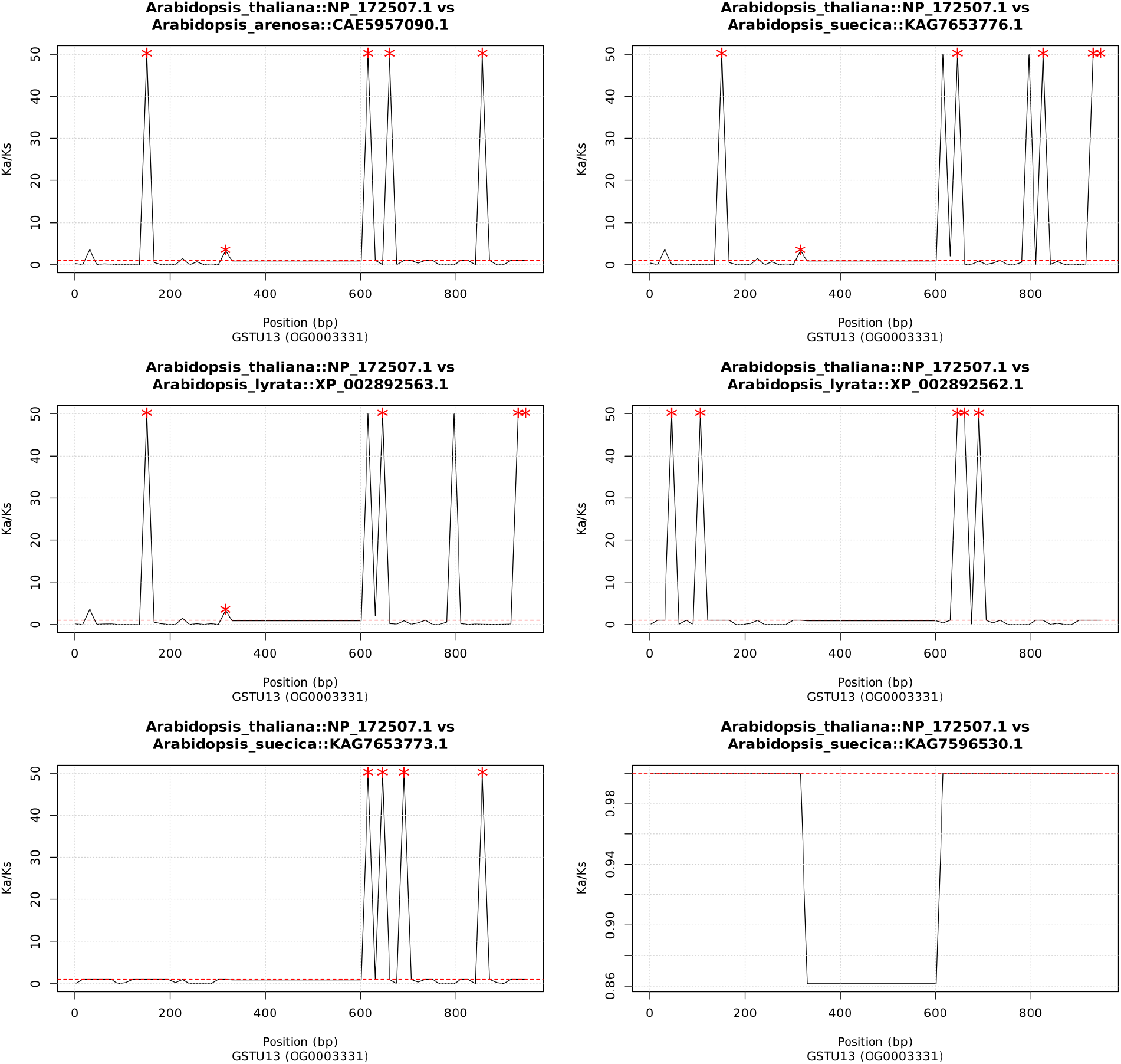
Ka/Ks (ratio of the number of nonsynonymous substitutions per non-synonymous site to the number of synonymous substitutions per synonymous site per unit time) across non-overlapping 15-bp sliding windows in GSTU13 (tau class glutathione transferase) gene between four Arabidopsis species. Red asterisks refer to windows with significant deviations from neutral expectations at α=0.1%.

