## Supplementary figures and images for "compare_genomes: a comparative genomics workflow to streamline the analysis of evolutionary divergence across genomes"

### Supplementary Figure 1

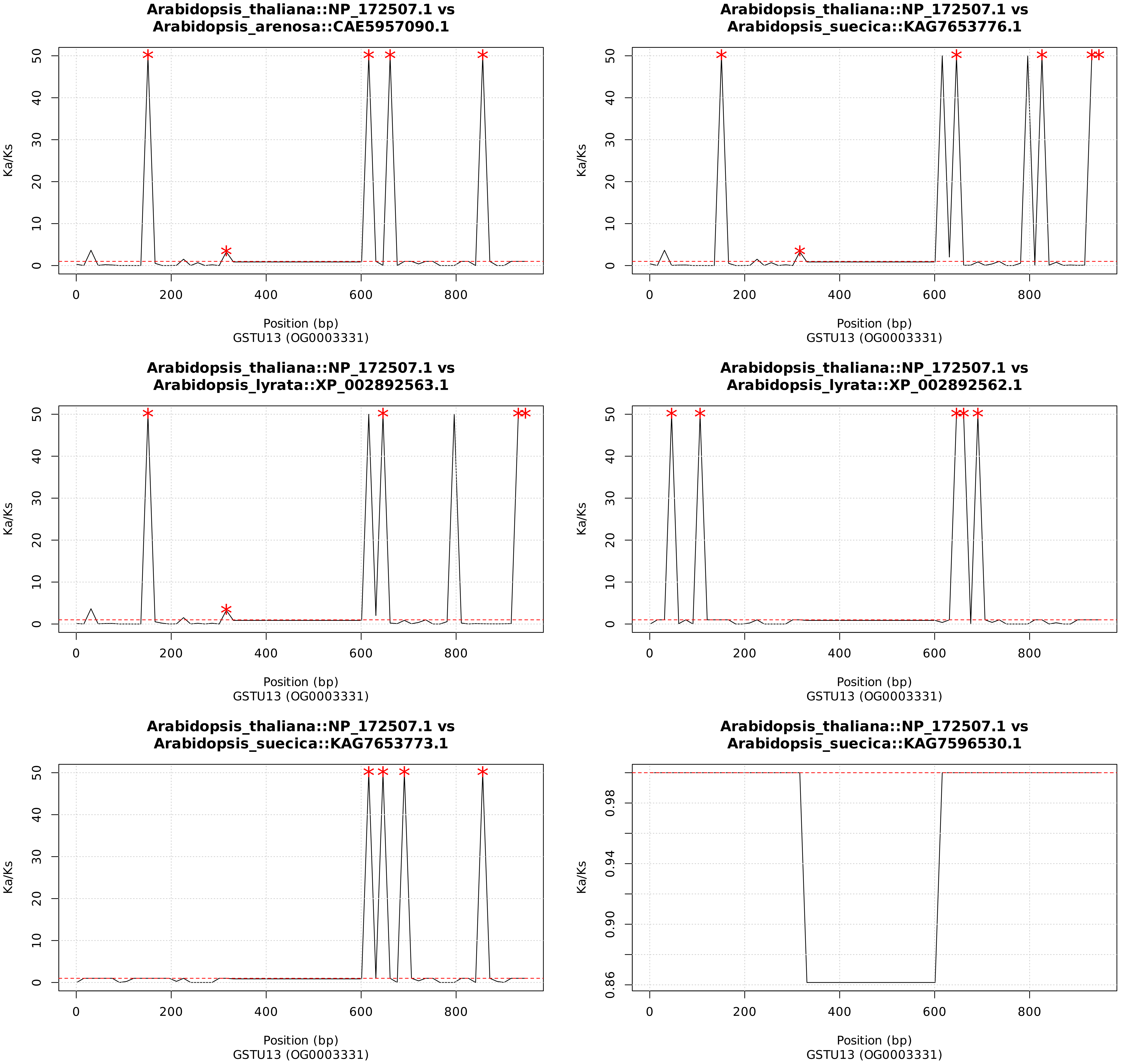
